# A large-volume sputum dry storage and transportation device for molecular and culture-based diagnosis of tuberculosis

**DOI:** 10.1101/2021.09.02.458812

**Authors:** Andrea Dsouza, Saylee Jangam, Vishwanath Naik, J. Manjula, Chandrasekhar B. Nair, Bhushan J. Toley

**Affiliations:** Department of Chemical Engineering, Indian Institute of Science, Bangalore, 560012, India; Centre for Biosystems Science and Engineering, Indian Institute of Science, Bangalore, 560012, India; Bigtec Labs, 2nd Floor, Golden Heights, 59th ‘C’ Cross, 4th M Block, Rajajinagar, Bengaluru, 560010, India

**Keywords:** Specimen stabilization, dry storage, dry blood spot cards, sputum transportation, tuberculosis diagnosis

## Abstract

Technologies for preservation of specimens in the absence of cold chains are essential for optimum utilization of existing laboratory services in the developing world. We present a prototype called specimen transportation tube (SPECTRA-tube) for the collection, exposure-free drying, ambient transportation, and liquid state recovery of large-volume (>1 mL) specimens. Specimens introduced into SPECTRA-tube are dried in glass fiber membranes, which are critical for efficient liquid-state sample recovery by rehydration and centrifugation. *Mycobacterium smegmatis (Msm)*-spiked mock sputum dried in native Standard 17 glass fiber was stable for molecular testing after 10-day storage at 45°C, and for culture testing after 10- and 5-day storage at 37°C and 45°C, respectively. Compatibility with human sputum storage was demonstrated by dry storing *Mycobacterium bovis*-spiked pooled human sputum in SPECTRA-tube for 5 days at room temperature followed by successful qPCR detection. By significantly increasing the volume of samples that can be transported in the dry state and enabling recovery of the entire sample in liquid state, SPECTRA-tube presents a potential universal solution for the preservation and transportation of liquid specimens.

---

Diagnostic services are essential to guiding treatment of many health conditions. In the developing world, diagnostic services are limited to well-equipped central laboratories in urban areas and the capability to conduct medical diagnostics rapidly drops in the remote regions. A large population that lives in remote areas, therefore, does not have access to state-of-the-art medical diagnostic facilities^1^. Central diagnostic laboratories in the developing world are well equipped to conduct high throughput and high-quality diagnostic testing but their potential is often underutilized^2^. An important reason for such underutilization is the absence of efficient cold chains for sample transportation resulting in poor quality of specimens. Specimens may often take several days to reach from sites of collection to the central laboratories and in the absence of proper stabilization they may be unfit for testing. Technologies for extended sample stabilization during transportation in the absence of cold chains will lead to better utilization and integration of the central laboratories into diagnostic workflows in the developing world^2^.

One approach to stabilizing specimens in the absence of cold chains is to dry them on a solid substrate, e.g. in dried blood spot (DBS) cards. DBS cards have been widely used for collection, preservation, transportation, and downstream molecular and immunological testing^2,3^. However, there has been little technological advance in specimen dry storage technology since the advent of DBS cards. DBS cards have several limitations: i) limited volumetric capacity (<100 μL per spot), ii) risk of contaminating the environment during specimen drying, iii) non-uniform sample distribution over the membrane, iv) the need to punch paper to retrieve sample which increases handling and biohazard, and v) poor efficiency of elution of the sample from paper^3^. Owing to these limitations, DBS cards (or current specimen dry storage technologies in general) have failed for applications where larger sample volumes may be needed.

An important application for dry storage of large specimen volumes is in the diagnosis of tuberculosis (TB) from sputum. TB, caused by *Mycobacterium tuberculosis* (*Mtb*), is the largest cause of deaths from a single infectious agent worldwide and claimed ~1.5 million lives in 2018, the majority of which were in developing nations^4^. Diagnostics is known to be a weak link in global TB control^5^. Effective stabilization of sputum from the site of collection to diagnostic laboratories, in the absence of cold chains, would significantly improve case finding and disease management. Because the concentration of *Mtb* in infected sputum may be as low as 100 bacteria per ml^6^, it becomes necessary to collect a large volume (>1 mL) for effective diagnosis. The primary requirements for sputum stabilization for TB diagnosis are: i) preserving *Mtb* nucleic acids for molecular testing, ii) preserving the viability of *Mtb* for culture testing, and iii) preventing the growth of non-tuberculous bacteria, which could grow at a significantly faster rate than *Mtb.* Research on stabilization of sputum has largely focused on transportation in a liquid state, e.g. using cetylpyridinium chloride (CPC)^7–9^ and ethanol^9^. PrimeStore MTM (Longhorn) is a commercial liquid sputum transport medium that lyses *Mtb* cells and preserves nucleic acids for molecular testing^10^. There have only been a few studies on dry sputum storage. Compared to transportation in liquid state, dry storage minimizes the risk of spillage during transportation. In 2001, one study reported dry storage of sputum on filter paper discs for culture and PCR tests^11^. Whatman FTA® DBS cards and GenoCard (Hain Life Sciences), which are commercial nucleic acid preservation cards, have been tested for dry sputum storage in a few studies^12–14^. A filter paper-based device to concentrate *Mtb* bacteria for sputum microscopy and molecular drug susceptibility testing was described by Tyagi et al^15^. TBSend – an end to end solution for collection and drying of sputum on filter paper and downstream molecular testing is being developed by Wobble Base Bioresearch^16^.

In this work, we present a prototype called SPECTRA-tube (specimen transportation tube) that enables spreading and rapid drying of a large volume of mock sputum over glass fiber membranes placed inside a centrifuge tube. The entire sample can be recovered in liquid state by rehydration and centrifugation within the same tube, obviating the need to punch the membrane for sample recovery. Recovery of the sample in liquid state renders it compatible with all routine laboratory diagnostic tests. We show that glass fiber membranes are enabling for sample recovery via centrifugation and outperform traditionally used cellulosic membranes. Using accelerated ageing, we demonstrate successful molecular and culture testing of *Mycobacteria* after long term dry storage in glass fiber membranes. By enabling exposure-free dry storage and efficient recovery of large specimen volumes, SPECTRA-tube represents a significant step ahead in specimen dry storage technology. In combination with a technology developed in our lab that enables efficient mixing of large fluid volumes with dried reagents in paper^17^, SPECTRA-tube promises to be a universal device for the stabilization of large specimen volumes.

## Results and Discussion

### Design and workflow of SPECTRA-tube

SPECTRA-tube consists of a stack of glass fiber membranes sandwiched in between two perforated acrylic layers (Fig. 1A). Multiple layers of partially cured PDMS (PDMS tape) are used as adhesive and spacers to secure the assembly and to prevent the glass fiber layers from collapsing, respectively. A 3D printed funnel feeds liquid sample to the glass fiber layers and is attached to one of the short edges of the assembly through a slot engraved into the acrylic layers. This assembly constitutes the sample cassette (Fig. 1A). The sample cassette is designed to fit snugly within a 50 mL Corning centrifuge tube (Fig. 1B). Two desiccant packs are placed on either side of the sample cassette for rapid drying of the sample contained within the glass fiber layers (Fig. 1B). Images of the sample cassette and the desiccant packs are shown in Figures 1C(i)–(ii). In this embodiment, the cassette consisted of two Standard 17 glass fiber layers (8 cm x 2 cm each). The desiccant packs were designed by folding pressure sensitive adhesive (PSA) tape into perforated pouches that were filled with color-indicating silica gel beads. A handle was designed into each pouch for ease of removal from SPECTRA-tube (Fig. 1C(ii)).

**Figure 1.**
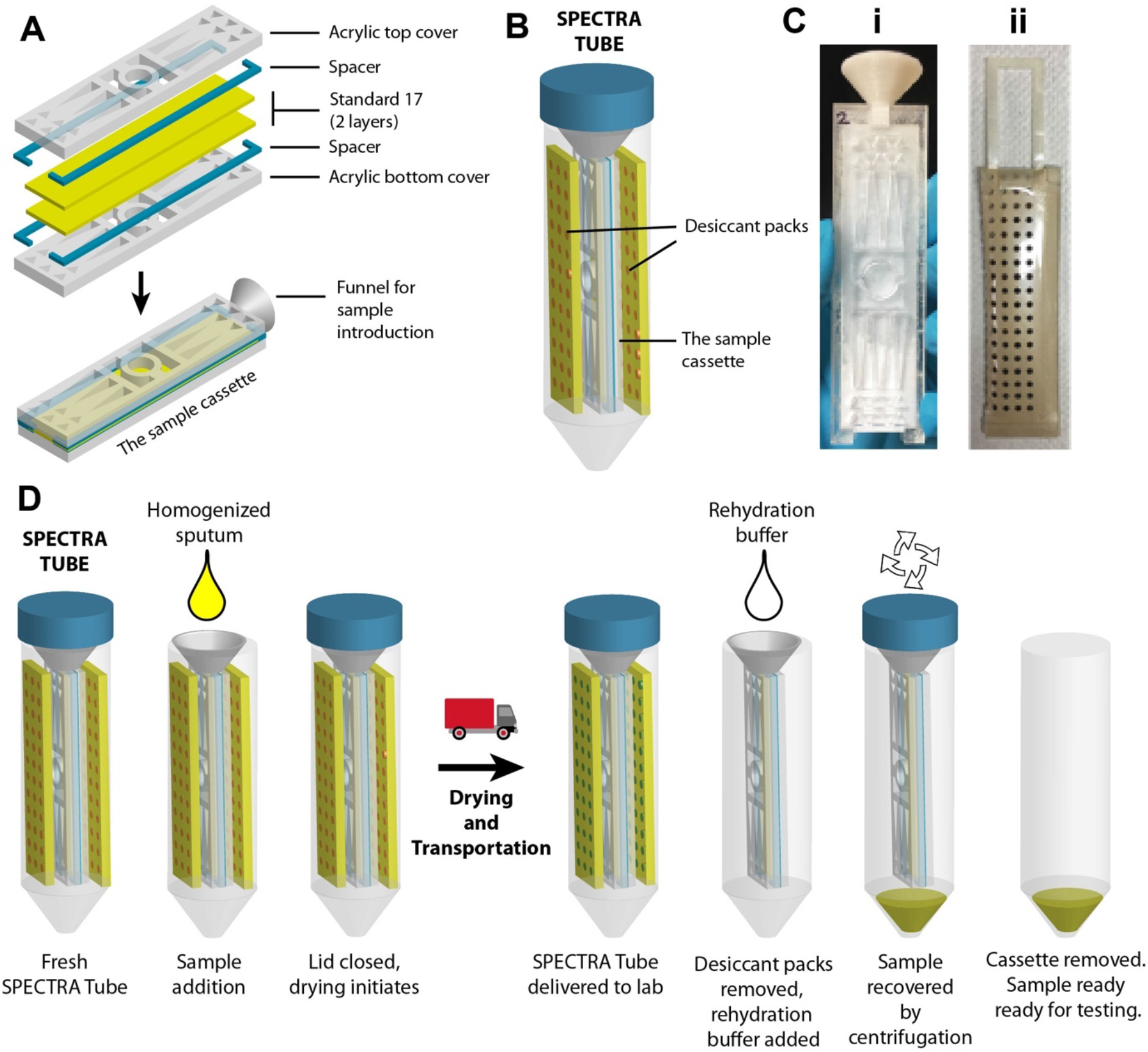
Design of SPECTRA-tube and workflow for sample collection and recovery. **A**. Part-by-part assembly of the SPECTRA-tube sample cassette. **B**. SPECTRA-tube consists of the sample cassette placed within a centrifuge tube surrounded by desiccant packs. **C**. Image of the sample cassette (i) and the desiccant pouch (ii). **D**. Workflow showing collection, drying, transportation, rehydration, and recovery (by centrifugation) of homogenized sputum.

The workflow for tuberculosis diagnosis from sputum collected remotely using SPECTRA-tube is shown in Fig. 1D. The sputum sample is first homogenized by mixing with a sputum homogenization solution, e.g. dithiothreitol (DTT) in ethanol^18^, and added into the SPECTRA-tube. The tube is capped to initiate drying and is now ready for transportation. After the sample reaches the diagnostic lab, the desiccant packs are removed, and a rehydration buffer is added to reconstitute the sample. The tube is capped and centrifuged to recover the reconstituted sample at the bottom of the tube. Recovery by centrifugation is critical because it enables recovering the entire sample, which may be necessary in case the target concentration in the sample is low. This is a marked improvement over previous sample dry storage techniques which are based on punching a part of the membrane for sample recovery.

### Choice of membrane for sample dry storage in SPECTRA-tube

Because sample recovery by centrifugation is central to the workflow of SPECTRA-tube, a membrane that maximized sample recovery by centrifugation had to be chosen. This was accomplished using a series of experiments that tested the recovery of different types of dried species from multiple types of membranes by rehydration and centrifugation. The three species that were dried and recovered were: i) a food coloring dye, ii) purified *Mycobacterium smegmatis* (*Msm*) genomic DNA in water, and iii) *Msm* cells in mock sputum. *Msm* was used as a BSL1-compatible surrogate for *Mtb*.

#### Recovery of dried food dye

As a preliminary examination, the efficiency of several membranes to release dried orange food coloring dye was quantified. Four types of membrane materials were chosen – a glass fiber membrane (Standard 17), a cellulosic membrane (Whatman Filter Paper 1), a nitrocellulose membrane (NCFF120HP), and a commercial nucleic acid storage membrane (FTA card). Square (1 cm^2^) pieces of each membrane were saturated with a dilute solution of the dye, dried for an hour in open air, rehydrated with water, and centrifuged to release fluid. Visual examination revealed that Standard 17 glass fiber was most effective at releasing dried orange dye (Fig. 2A). There was no trace of the dye visible in Standard 17 while all other membranes exhibited different intensities of orange color after centrifugation. Filter paper 1 and FTA® card, both cellulosic membranes, showed maximum retention of the orange dye. Because glass fiber membranes outperformed membranes made from other materials, 4 types of glass fiber membranes were then compared: Standard 17, Standard 14, GF/DVA, and Fusion 5. Among these, Standard 17 exhibited the lowest retention visually (Fig. 2B).

**Figure 2.**
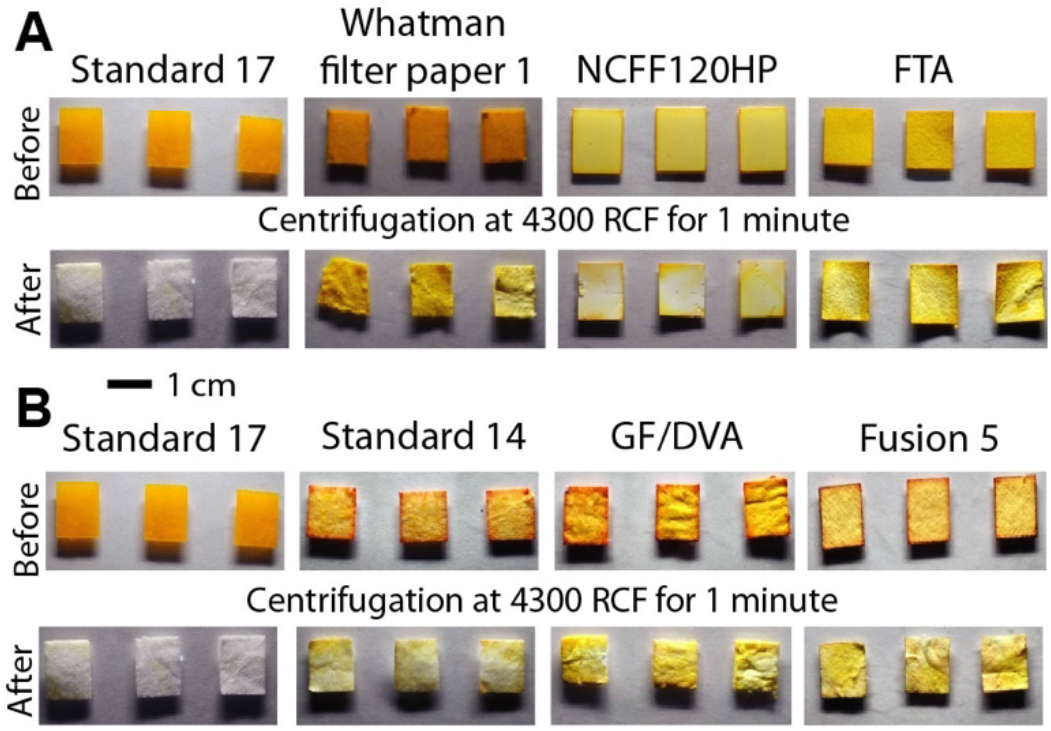
Recovery of dried orange dye from different membranes. **A**. Images of glass fiber Standard 17, Whatman filter paper 1 (cellulose), nitrocellulose FF120HP, and Whatman FTA® membranes, before and after recovery of dried orange dye. **B**. After recovery, Standard 17 had a lower mean color intensity compared to filter paper 1 (*; P < 0.05; N = 3), NCFF120HP (*; P < 0.05; N = 3), and FTA® card (**; P < 0.001; N = 3). **C**. Images of various glass fiber membranes before and after recovery of dried orange dye. **D**. After recovery, Standard 17 had a lower mean color intensity compared to Standard 14 (**; P < 0.01; N = 3), GF/DVA (***; P < 0.001; N = 3), and Fusion 5 (**P < 0.01; N = 3).

#### Recovery of dried genomic DNA

The efficiency of releasing dried purified *Msm* genomic DNA from different membranes was tested next. The workflow for sample drying, rehydration, and recovery was as shown in Fig. 3A. A solution containing the DNA was introduced into square (1 cm^2^) membranes of each material, dried in a petri dish containing desiccant, and stored for 24 h. Post storage, membranes were rehydrated, fluid was recovered by centrifugation, and qPCR was used for DNA quantification. First, three types of membranes: glass fiber Standard 17, cellulosic filter paper 1, and nitrocellulose NCFF120HP were compared. A genomic DNA solution that was never dried was used as a control. The concentration of DNA in the sample recovered from Standard 17 was not statistically different from controls (N=3; P = 0.84; Fig. 3B), which shows that there were no significant losses in recovery. Concentrations of DNA in solutions recovered from filter paper 1 and NCFF120HP were significantly lower than the control (***; P < 0.001; N=5; Fig. 3B). In a separate experiment, the efficiency of releasing dried genomic DNA was tested across different glass fiber membranes – Standard 17, Standard 14, GF/DVA, and Fusion 5. The concentration of DNA recovered from Standard 17 was statistically higher than Standard 14 (*; P < 0.05; N=3), GF/DVA (**; P < 0.01; N=3), and Fusion 5 (*; P < 0.05; N=3) (Fig. 3C). Thus, even among the glass fiber membranes tested, Standard 17 was most efficient at releasing stored DNA.

**Figure 3.**
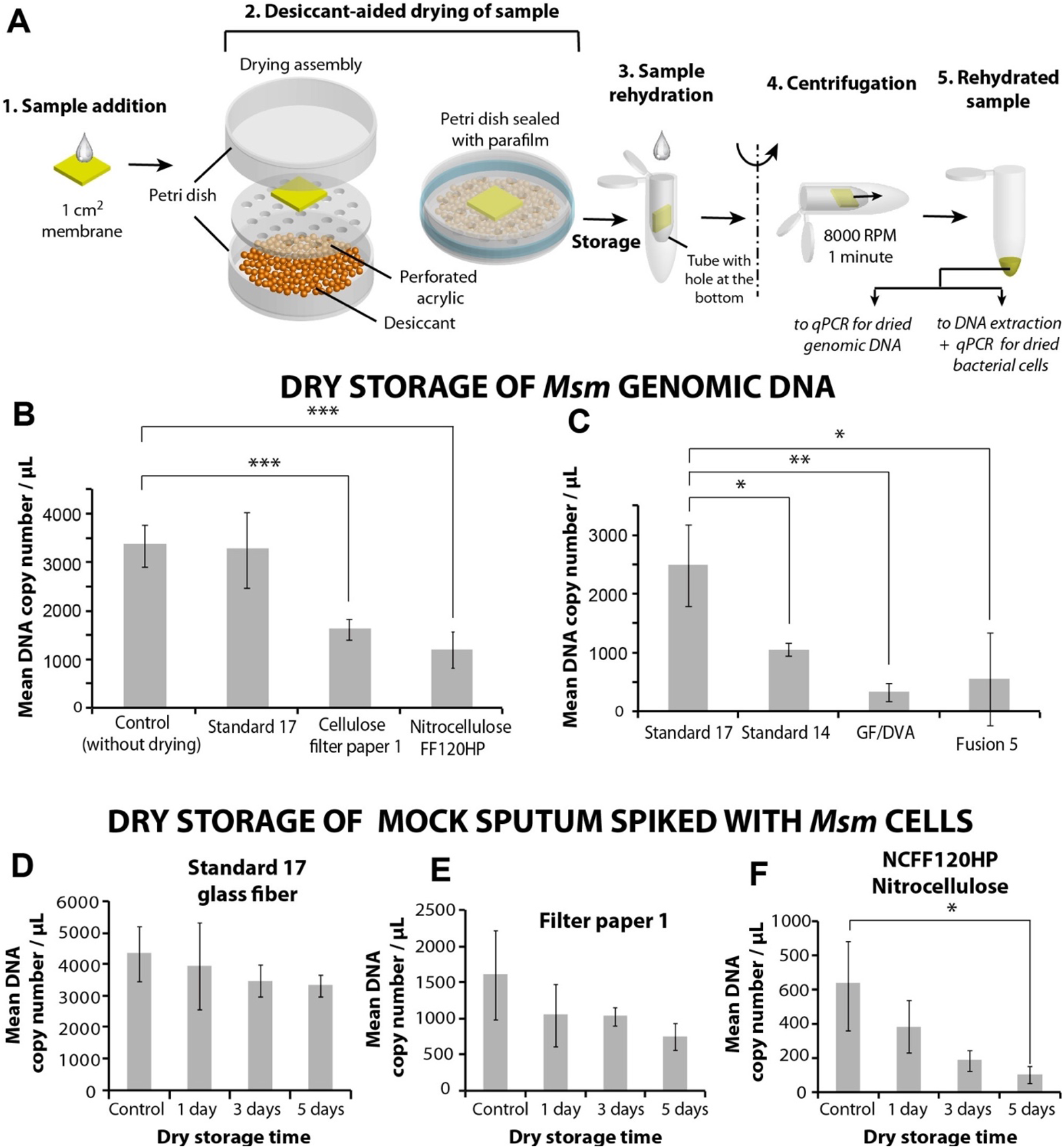
Recovery of dried genomic DNA and *Msm* bacteria from different membranes. **A**. Schematic representation of the workflow for sample drying and recovery. **B**. Comparison of recovered DNA concentrations across membranes made of different materials. Compared to control, recovered DNA concentrations were significantly lower for filter paper 1 (***; P < 0.001; N=3) and NCFF120HP (***; P < 0.001; N=3), but not for Standard 17. **C**. Comparison of recovered DNA concentrations across glass fiber membranes. Compared to Standard 17, recovered DNA concentration was significantly lower in Standard 14 (*; P < 0.05; N=3), GF/DVA (**; P < 0.01; N=3), and Fusion 5 (*; P < 0.05; N=3). All error bars represent standard deviations. **D-F**. Dry storage of Msm-spiked mock sputum for 5 days in Standard 17 (**D**), filter paper 1 (**E**), and NCFF120HP (**F**). The loss in quantifiable DNA was least in Standard 17.

#### Recovery of dried Msm bacterial cells in mock sputum

Mock sputum (viscosity 6.9 cP) containing *Msm* bacteria (10^3^ CFU/ml) was dried in square (1 cm^2^) membranes of Standard 17, Whatman filter paper 1, and NCFF120HP, stored at room temperature, and recovered after 1, 3, and 5 days using the workflow shown in Fig. 3A. Fresh undried mock sputum was used as a control. DNA was first extracted from the samples using a QIAamp DNA mini kit followed by qPCR for quantification. For all three membranes, there was a trend of decreasing DNA concentration with increasing days of storage, but the reduction was minimum in Standard 17 (Fig. 3D–F). There were no statistically significant losses in quantifiable DNA compared to the control after 1, 3, and 5 days of storage in Standard 17 (Fig. 3D). Similarly, for Whatman filter paper 1, the concentrations compared to control after 1, 3, and 5 days storage were not statistically significant (Fig. 3E). For NCFF120HP, DNA concentration after 1 and 3 days was not significantly different, but after 5 days was significantly different compared to control (*; P<0.05; N=3; Fig. 3F). The percentage losses (compared to control) in mean DNA concentration after 5 days storage in Standard 17, filter paper 1, and NCFF120HP were 23%, 53%, and 92%, respectively (Fig. 3D–F). These preliminary studies indicated that Standard 17 glass fiber membrane would be most suitable for specimen dry-storage and recovery by centrifugation.

### Accelerated ageing studies for molecular testing

The feasibility of conducting molecular diagnostic tests from *Msm*-spiked mock sputum samples stored long-term in Standard 17 membranes was tested next using an identical method. Samples were stored at RT, 37°C, and 45°C and recovered after 1, 5, and 10 days, followed by DNA extraction and qPCR quantification. Despite losses in quantifiable DNA concentration, samples stored under all conditions contained amplifiable DNA (Fig. 4), demonstrating successful dry storage of mock sputum for 10 days at 45°C (equivalent to 40 days at RT, assuming doubling of reaction rate for every 10°C rise, according to Arrhenius equation). Recovered DNA concentration followed a decreasing trend with increasing storage time at all temperatures (Fig. 4). Compared to control, the losses were significant for the following conditions: RT day 10; 37°C days 5 and 10; and 45°C days 5 and 10 (*; P<0.05; Fig. 4). For 10-day storage, the percentage losses in mean DNA concentration at RT, 37°C, and 45°C were 73.8%, 88.1%, and 92.2%, respectively. Although these losses appear to be large when expressed as percentages, qPCR assays usually only concern themselves with *Ct* values. The maximum loss (92.2%) in detectable DNA concentration for 10 days storage at 45°C corresponds to a 12.6-fold reduction, which corresponds to an increase in qPCR *Ct* value of only 3.7 cycles (log212.6), assuming ideal qPCR conditions. Thus, the losses in quantifiable DNA even for 10 days storage at 45°C are not substantial. Note that because these qPCR assays were based on intercalating dyes, non-specific PCR amplification products could contribute to fluorescence signals and report falsely elevated concentrations of target DNA. To rule this out, melt curve analysis as well as gel electrophoresis was performed on all PCR products. All melt curves featured a single peak (Supplementary Figure S1) and all gels featured a single target band (Supplementary Figure S2), which confirmed the absence of non-specific amplification products.

**Figure 4.**
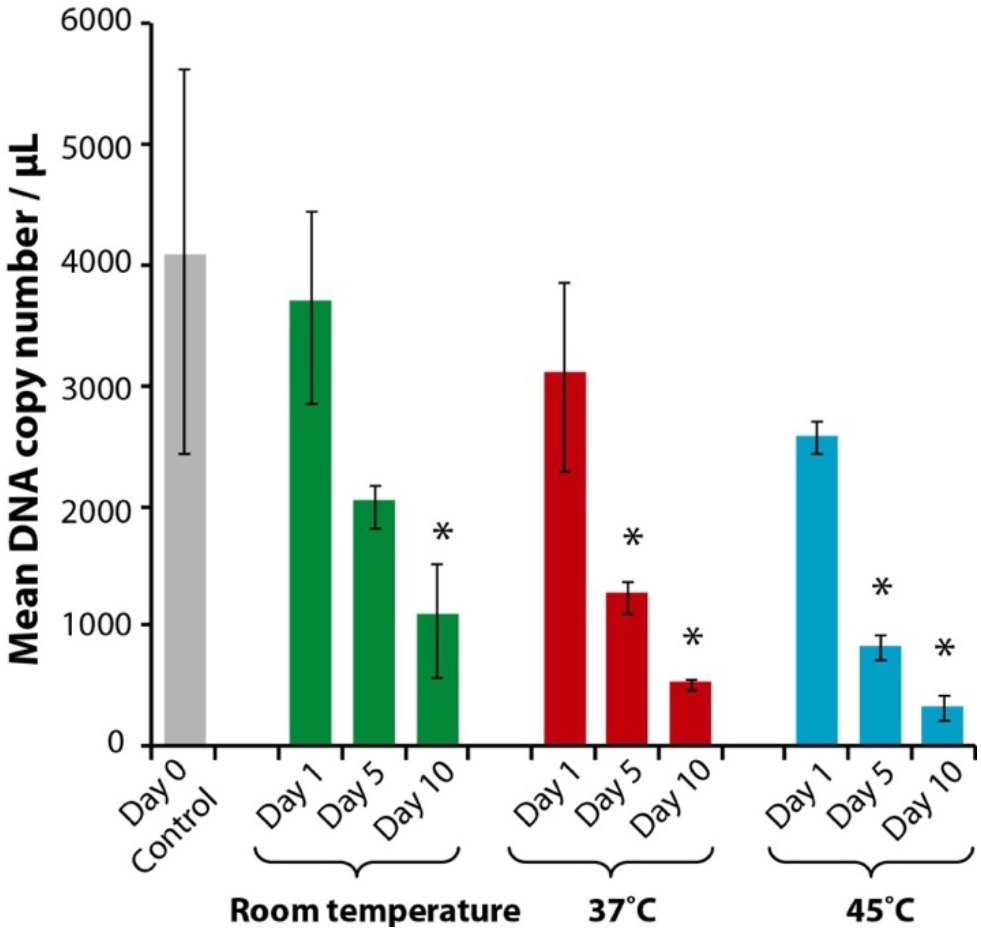
Accelerated ageing studies for molecular detection. Quantitation of *Msm* DNA concentration in mock sputum dried in Standard 17 and stored at room temperature, 37°C, and 45°C. All statistical comparisons are made against the control which was a sample that was never dry stored (*; P<0.05; N=3).

### Accelerated ageing studies for culture testing

The feasibility of conducting culture tests from *Msm*-spiked mock sputum samples stored long-term in Standard 17 membranes was tested next. Samples were stored at RT, 37°C, and 45°C and recovered at days 1, 5, and 10 for plating. Colonies formed were treated with a nucleic acid stain EtBr. EtBr is taken up by non-*Mycobacterial* cells due to the presence of *LrfA* transporter in their cell membrane producing fluorescent colonies (Fig. 5Ai). *Mycobacterial* cells lack the transporter and thus do not uptake EtBr and do not produce fluorescence (Fig. 5Aii). This test was used to enquire whether colonies forming from recovered samples were constituted of *Mycobacteria*. When stored at RT and 37°C, large number of colonies were observed for 1, 5, and 10 days of storage (Fig. 5B). When stored at 45°C, colonies were clearly visible after 1 day of storage, but the colony count dropped for 5 and 10 days of storage (Fig. 5B). All colonies produced in these experiments were non-fluorescent, indicating that they were *Msm* colonies (Fig. 5B and Supplementary Figure S3). Colony culture PCR performed from a few colonies produced a product of the expected band size (185 bp), which confirmed *Msm* colonies (Fig. 5C and Supplementary Figure S4). A plot of colony counts for each storage condition is shown in Fig. 5D. Despite the drop in the number of colonies, viable bacteria persisted under all conditions, except at 45°C Day 10, for which only 1 out of the 3 replicate plates produced colonies (Supplementary Figure S3). Based on the above data, Standard 17 was finalized as the membrane for SPECTRA-tube.

**Figure 5.**
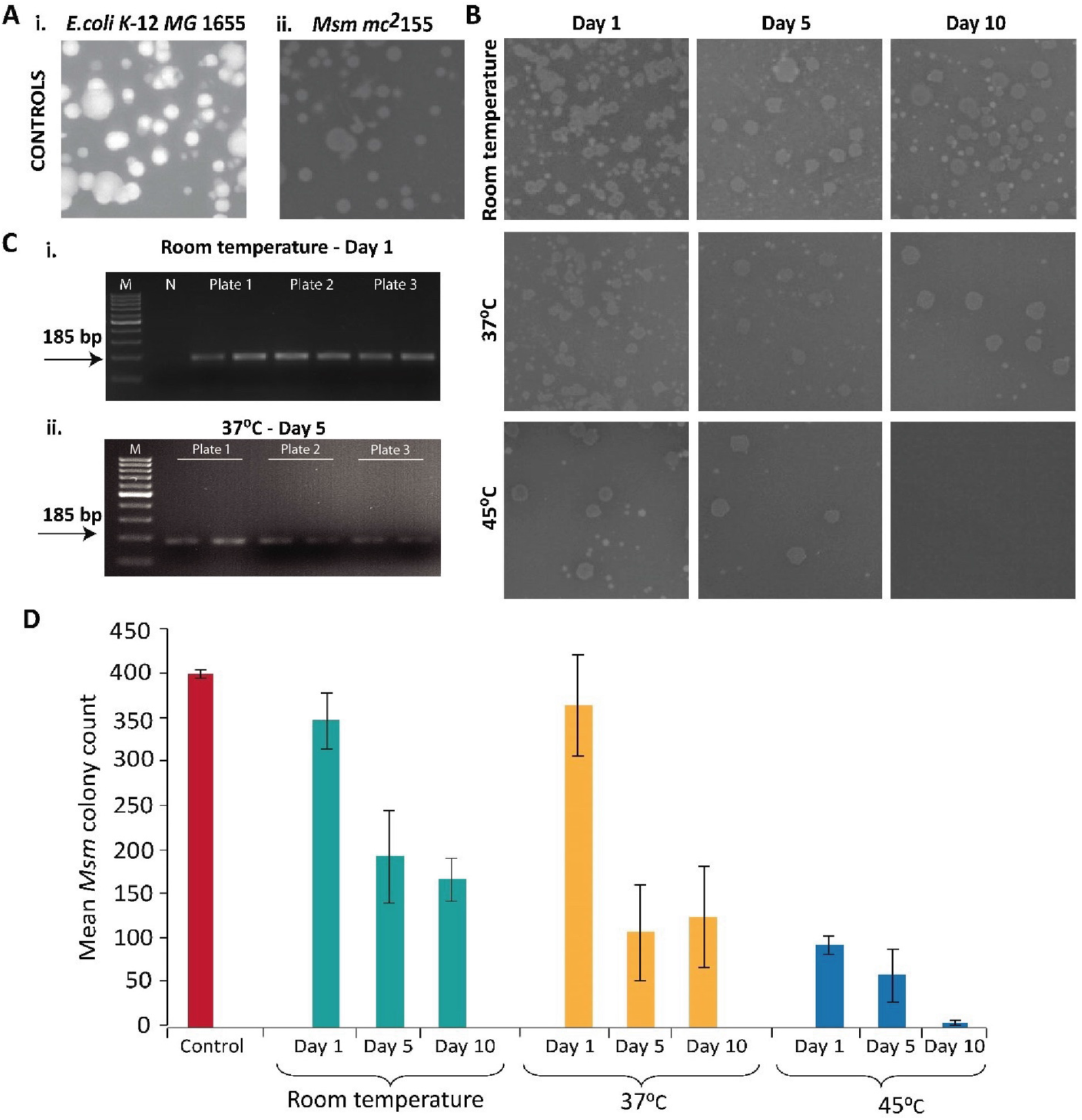
*Msm* viability in dried mock sputum samples. **A.** Fluorescent images of control cultures showing EtBr uptake by model non-*Mycobacterial* cells, *E.coli* (i) and lack of EtBr uptake by *Msm* cells (ii) **B.** Images of agar plates showing *Msm* colony growth from samples stored under different conditions. **C.** Agarose gel electrophoresis images of products of colony-culture PCR for a few representative storage conditions showing bands at 185 bp, confirming growth of *Msm*. M: marker; N: negative control. **D.** Plot of *Msm* colony counts obtained from samples stored under different conditions. Error bars represent standard deviations (N=3).

### Sputum drying rate in SPECTRA-tube

After Standard 17 was finalized as the membrane for SPECTRA-tube, the drying rate of mock sputum in SPECTRA-tubes was measured. Mock sputum volumes of 0.6, 1.2, and 1.8 mL were added to SPECTRA-tube sample cassettes containing 1, 2, and 3 layers of Standard 17, respectively. Cassettes were removed every 2 hours and their weight measured. A different SPECTRA-tube was used for every time point to avoid alteration in drying rate by opening of the tube during weight measurement. Each SPECTRA-tube contained two desiccant pouches containing ~5.4 g silica gel (2.7 g each). Assuming that the water adsorption capacity of silica gel is ~30% of its weight^19^, the theoretical capacity for water adsorption in each tube was ~1.62 g. For 1-layer devices, 97.7% of 0.58 g added sputum dried in 4 hours; for 2-layer devices, 96.7% of 1.16 g added sputum dried in 8 hours; for 3-layer devices, only 77.2% of 1.74 g added sputum dried in 8 hours because the added weight was more than the theoretical adsorption capacity of the silica gel (1.62 g). Subsequent experiments were therefore only conducted using 1- or 2-layer cassettes in which sputum could be efficiently dried within 8 hours.

### Optimization of sample recovery from SPECTRA-tube

Dried samples were recovered from SPECTRA-tube by addition of water (equal volume to that of added specimen) followed by centrifugation. Centrifugation speeds and times were optimized (Supplementary Table 1) for maximization of the weight of rehydrated sample from SPECTRA-Tube devices and finalized as 6,037 RCF for 3 min followed by 16,770 RCF for 3 min.

### Comparison of FTA card and SPECTRA-tube

Nucleic acid recovery from sputum stored in a 1-layer SPECTRA-tube was directly compared to FTA cards using a workflow shown in Fig. 7A. *Msm*-spiked (10^3^ CFU/ml) mock sputum was introduced into each device and the volumes were chosen to match the volumetric capacities – 640 μl for the 1-layer SPECTRA-tube and 125 μl for a 3.5 cm diameter disc of FTA card. After drying at RT and storage for 24 hours, samples were recovered from the FTA card using a 5 mm punch and recovered from SPECTRA-tube using centrifugation. All samples underwent nucleic acid extraction followed by PCR. Distinct 185 bp bands were visible on gels from samples stored in SPECTRA-tube (Fig. 7B). However, no amplifiable DNA was present in any sample stored in FTA cards, evident from the absence of (or extremely low intensity of) the 185 bp band (Fig. 7C). The improved ability of SPECTRA-tube for storage and detection of *Msm* stems from its larger volumetric capacity and the ability to efficiently recover the entire sample using centrifugation. FTA cards, on the other hand, have a smaller volumetric capacity and enable recovering only a small part of the sample by punching. These results show that SPECTRA-tube outperforms commercially available nucleic acid storage membranes for sputum dry storage.

**Figure 6.**
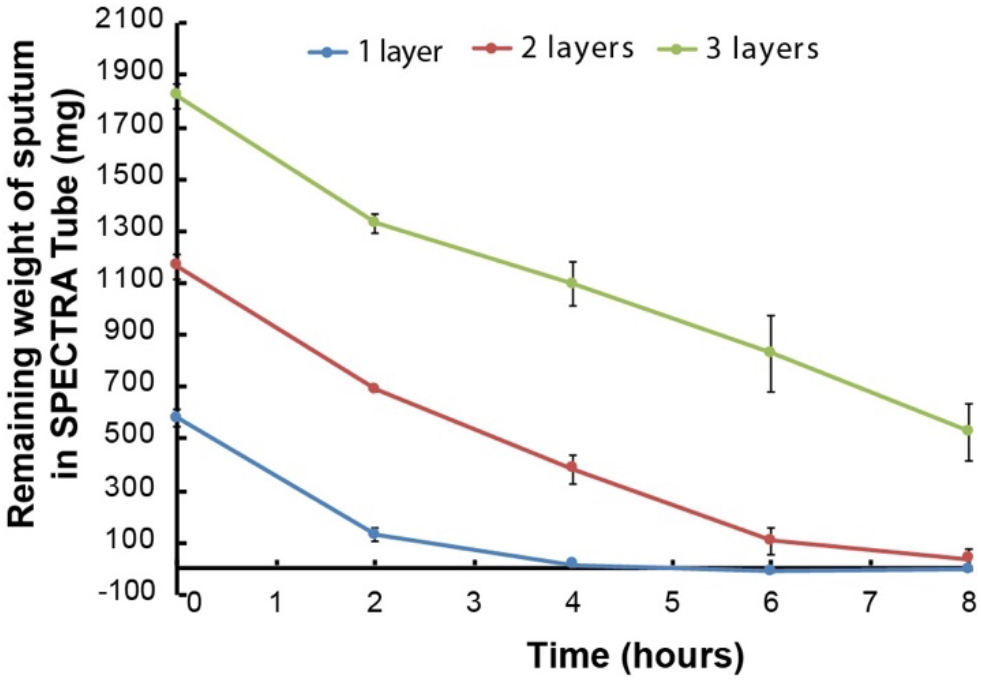
Drying rate of mock sputum in SPECTRA-tube. Weight of mock sputum remaining in the glass fiber layers of SPECTRA-tube during the process of drying, for 1-layer, 2-layer, and 3-layer SPECTRA-tube devices. Error bars represent standard deviations (N=3).

**Figure 7.**
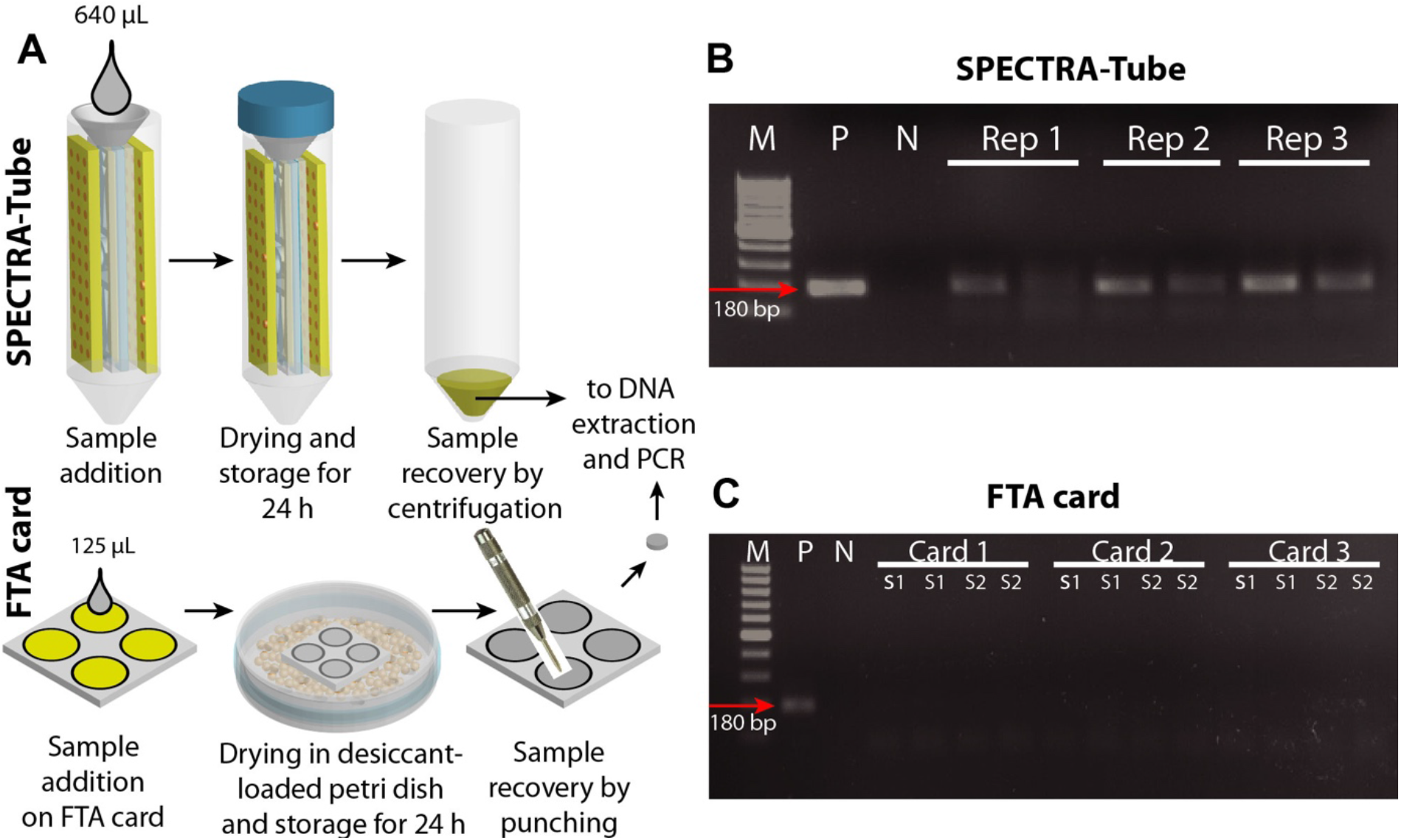
Comparison of sputum recovery from SPECTRA-tube and FTA cards. **A.** Workflow of mock sputum dry-storage and recovery from SPECTRA-tube and FTA cards. **B-C.** Results of agarose gel electrophoresis. Sputum recovered from SPECTRA-tube showed 185 bp bands indicating presence of *Msm* DNA (B). Absence of the 185 bp band indicates that *Msm* DNA was not detectable in sputum recovered from FTA cards (C). M: marker; P: positive control; N: negative control; Rep 1, Rep 2, Rep 3: SPECTRA-tube replicates; two lanes for each Rep are PCR duplicates. Card 1, Card 2, Card 3: FTA card replicates; S1, S2: Two different 5 mm discs punched from a single FTA card; two lanes for S1 and S2 show PCR duplicates.

### Dry storage of human sputum in SPECTRA-tube

Pooled human sputum was prepared at Bigtec Labs as a mixture of multiple discarded sputum samples from patients that tested negative for TB, and spiked with *Mycobacterium bovis*. Multiple 1.2 mL pooled sputum samples were introduced into 2-layer SPECTRA-tubes and stored for 5 days. Samples were rehydrated with 1.2 mL DI water and recovered by centrifugation followed by DNA extraction and qPCR. The *Ct* values for controls (fresh samples) and samples stored for 5 days in SPECTRA-tubes were 27.8 ± 0.09 and 29.6 ± 0.16 (N=3), respectively, with a mean increase in *Ct* of 1.8 cycles, which indicates a 3.4X loss in detectable DNA concentration. Despite the modest loss, this result demonstrates the application of SPECTRA-tube for human sputum storage.

SPECTRA-tube presents a paradigm shift in specimen dry storage technologies. It enables i) drying of a significantly larger sample volume than has traditionally been possible with DBS card-like technologies, ii) rapid drying of samples in enclosed tubes avoiding sample exposure, and iii) recovery of the entire sample in liquid state by centrifugation (as opposed to by punching small regions of a membrane). SPECTRA-tube is a versatile platform that can be modified for different types of samples and applications. The sample storage capacity of SPECTRA-tube is limited only by the amount of desiccant. The prototype presented here accommodates ~5.4 g silica gel, but further design changes could be made to increase desiccant loading capacity. The number of Standard 17 layers could be increased to increase fluidic capacity. Note that unlike commercial nucleic acid storage membranes like FTA cards, nucleic acid stabilizing reagents were not used in SPECTRA-tube. In theory, DNA could be stabilized for significantly longer periods of time if *Mycobacterial* cells were lysed chemically and all proteins in the sample were denatured (as is done in FTA cards). However, lysis renders the sample incompatible with culture testing. Sputum dry storage in Standard 17 membranes in SPECTRA-tube may thus be the optimal solution that enables a full sputum workout for tuberculosis diagnosis, i.e. molecular and culture testing. However, for applications not requiring culture testing, nucleic acid stabilization reagents may easily be incorporated into SPECTRA-tube. To enable this, our group has developed a novel stacked paper design that enables efficient mixing of large fluid volumes with dried reagents stored in paper membranes, and demonstrated its application in sputum decontamination^17^.

This work also sheds light on the suitability of glass fiber membranes for specimen dry storage. Till date, most attempts to dry liquid specimens had been made using cellulose-based membranes. Compared to cellulose, glass fibers have larger pore sizes, lower capillary pressure, less resistance to flow, and higher volumetric capacity^20^. Low resistance to flow is an important feature for wicking viscous fluids, and low capillary pressure is ideal for sample recovery using centrifugation. When a biological fluid is introduced into a cellulose-based paper membrane, the components of the fluid are absorbed into the cellulose fibers; in contrast, in glass fibers, the components dry out as films^21^. Glass fiber membranes also have higher mechanical strength compared to cellulose-based membranes^21,22^.

The total cost of materials required to build a 2-layer SPECTRA-Tube (that holds 1.2 mL sputum) is ~$1.38 (~INR 101; Supplementary Table 2). Out of this, the major cost is that of the Corning centrifuge tube (INR 31) and the Standard 17 membranes (INR 35), which together account for ~65% of the cost. The cost must be further reduced for widespread use in the developing world. The cost of the centrifuge tube can easily be reduced by acquiring it locally. Replacement of Standard 17 with alternative materials acquired locally and a revamp of the cassette design to make it monolithic would be a subject of further research in our group. Additionally, the current prototype will be validated for TB diagnosis from dried sputum using a set of clinical samples as a part of our future work.

### Outlook

SPECTRA-tube is a new specimen dry storage technology that overcomes many limitations of existing specimen dry storage technologies, e.g. limited specimen volumes, sample drying in open air, and recovery of only a small fraction of the collected sample by punching. For TB diagnosis, it enables dry storage of sputum followed by both molecular and culture testing; the latter is not possible with any current sample stabilization technology. While demonstrated here for TB diagnosis from sputum, SPECTRA-tube is versatile and could be used as a universal platform for the dry storage of any kind of liquid specimen. In the developing world, SPECTRA-tube could improve the quality of medical diagnostic testing by bringing better quality specimens to the central laboratory. In the developed world, adaptations of SPECTRA-tube may be used for at-home collection and stabilization of different types of samples like urine and saliva for shipping to the diagnostic laboratory. Overall, SPECTRA-tube promises to increase the quality and penetration of medical diagnostic testing by enabling the safe collection, preservation, and transportation of larger volumes of liquid specimens than what has been possible till date.

## Experimental Section

### Membrane materials

Whatman nitrocellulose membrane NC FF120HP, glass fiber membranes Standard 17, Standard 14, GF/DVA, and Fusion 5 were procured from Wipro GE Healthcare Pvt. Ltd. (Bengaluru, India). Whatman FTA classic card was acquired from Sigma-Aldrich (WHAWB120305). All membranes were cut using a 50W CO2 laser cutter (VLS 3.60; Universal Laser Systems, Scottsdale, AZ).

### Fabrication and assembly of SPECTRA-tube

All parts of the device were designed using AutoCAD 2017 (Autodesk) and cut using a VLS 3.60 CO2 laser cutter. For the sample cassette, 2.6 mm acrylic sheets were used to sandwich multiple layers of 80 x 20 mm standard-17 glass fiber. PDMS tape (Arclad IS-7876) for securing the cassette was acquired from Adhesives Research, Glen Rock, PA. PSA tape for fabricating desiccant packs was acquired from 3M (product number 3791). Orange silica gel beads that turned green after adsorbing moisture, used in desiccant packs, were acquired from Cilicant, Pune, India. The funnel was designed using Autodesk Fusion 360 software and 3D printed in acetyl butadiene styrene (ABS) (Tesseract, Mumbai, India) using the Accucraft i250 D 3D printer (Divide by Zero, Mumbai, India).

### Bacterial culture and DNA extraction

*Mycobacterium smegmatis mc^2^*155 (*Msm*) was obtained as a kind gift from Prof. Deepak Saini (Indian Institute of Science, Bangalore). *Msm* was routinely grown in tryptone soya broth with 0.05% Tween 80 for 24 hours at 37°C, 180 RPM (primary culture), followed by growth in Middlebrook 7H9 broth with 2% glucose for 24 hours at 37°C, 180 RPM (secondary culture). Genomic DNA was extracted and purified using QIAamp DNA mini kit according to manufacturer’s protocol.

### Preparation of mock sputum and viscosity measurement

Mock sputum was prepared as a solution of 1.8% w/v methylcellulose and 10% egg yolk emulsion in sterile distilled water^17^. The viscosity of mock sputum was measured using a cone and plate rheometer MCR 301 (Anton Paar, Graz, Austria). Mock sputum was then diluted 3X in DI water, as carried out by the GeneXpert MTB/RIF test, to mimic homogenized sputum. All references to mock sputum in the described experiments refer to this 3X diluted solution.

### Dry storage and recovery experiments for comparison between membranes

Different volumes of either i) dilute food coloring dye solution, ii) purified *Msm* genomic DNA (10^3^ copies/μl) in water, or iii) *Msm*-spiked mock sputum (10^3^ CFU/ml of sputum) were introduced into 1cm^2^ square membranes until fully saturated (40 μl for Standard 17; 10 μl for Whatman filter paper 1 and Nitrocellulose FF120HP; 25 μl for FTA; 38 μl for Standard 14; 98 μl for GF/DVA; 52 μl for Fusion 5) and allowed to dry at room temperature. For orange dye experiments, drying was conducted in open air and for the other two samples, it was conducted in sealed petri dishes (Fig. 3A). For sample recovery, the membranes were rehydrated with equal volumes of water and centrifuged at 4300 RCF (8000 RPM; rotor radius = 6 cm) for 1 minute.

### Quantification of nucleic acids

DNA was quantified using qPCR performed in an Applied Biosystems QuantStudio 3 with Takara TB Green Premix Ex Taq II (Tli Rnase H Plus). Primers were designed for the *sigA* gene – forward: 5’-GACTCTTCCTCGTCCCACAC-3’ and reverse: 5’-GAAGACACCGACCTGGAACT-3’ to produce a 185 bp product. For *Msm*-spiked mock sputum, DNA was first extracted from each recovered sample using QIAamp DNA mini kit before qPCR. For each recovered sample, purified DNA was eluted into a final volume of 200 μl and 1 μL from that was used for qPCR. Nucleic acid extraction step was not performed for the case when pre-purified nucleic acids were dried in paper membranes. The qPCR reaction consisted of 15 μL master mix, 0.3 μL forward and reverse primers each, 1 μL target nucleic acids, and sterile water to adjust the total volume to 30 μL. In all cases, it was ensured that the qPCR efficiency was in between 90 to 100%. All statistical comparisons in this work were made using two-tail Student’s t-tests performed in Microsoft Excel.

### Accelerated ageing studies of dried mock sputum

Two types of accelerated ageing studies were conducted: i) for nucleic acid quantification from recovered samples, and ii) for determination of *Msm* viability from recovered samples. All experiments were performed on 1cm^2^ Standard 17 membranes which were autoclaved at 121?C, 15 psi pressure for 30 mins to ensure that the membranes were sterile. After autoclaving the membranes were further UV-sterilized for 10 minutes and stored in sterile airtight containers before use. *Msm*-spiked mock sputum (10^3^ CFU/ml) was introduced into the membranes on Day 0, stored at RT, 37°C, and 45°C, and recovered after 1, 5, and 10 days. Nucleic acids were quantified as described above. *Msm* viability was quantified using the following procedure. Recovered samples were serially diluted 10X and each sample was decontaminated with 1% NaOH. After decontamination, the samples were plated on Middlebrook 7H9 agar (N=3 plates for each sample) and incubated at 37°C for 48 hours before colony counting. Two techniques were used to confirm that colonies formed comprised of *Msm* bacteria: i) Ethidium bromide (EtBr) fluorescence staining, and ii) colony culture PCR. For EtBr assay, 500 μL of 0.1mg/ml EtBr in sterile water was introduced in each agar plate after colony formation, followed by 5 min incubation at RT and fluorescence imaging in a gel doc system (UVITech Essential V6). Colony-culture PCR was performed by picking 3 colonies from each agar plate and suspending them in tubes containing 1mL Middlebrook 7H9 media. The tubes were incubated at 37°C for 36 h followed by DNA extraction, PCR amplification of *sigA* gene, and agarose gel electrophoresis.

### Dry storage of pooled human sputum

SPECTRA-tubes were fabricated at IISc and shipped to Bigtec Labs where they were used for dry storage of pooled sputum. After recovery of dried sputum, DNA was extracted using the Trueprep Auto extraction system (Molbio Diagnostics, Goa, India) and amplified using an Applied Biosystems 7500 real time PCR machine. The *Mycobacterium tuberculosis* PCR assay amplified the *nrdB* gene and consisted of a reaction buffer, 2 units Taq polymerase, 400 nM dNTPs each, 0.4 μM forward and reverse primers, 0.1 μM probe, 3 mM MgCl2, 4 μL template, and water to adjust the final reaction volume to 10 μL.

## Acknowledgements

This work was supported by a Grand Challenges Exploration award from the Bill & Melinda Gates Foundation (OPP1182249), an extramural research grant from the Science and Engineering Research Board, India (EMR/2016/006029), an Innovative Young Biotechnologist award from the Department of Biotechnology, India (BT/010/IYBA/2016/07), a Grand Challenges Exploration-India award from BIRAC-India, and by the Saroj Poddar Foundation in the form of a Young Investigator award.

## Author Contribution Statement

B.JT contributed towards conception and design of the work and acquired funding. SJ designed and optimized the SPECTRA-tube prototype. AD conducted experiments to determine the optimal membrane for SPECTRA tube, designed the *Msm* qPCR assay, and conducted all molecular biology and microbiology experiments. CBN contributed towards design of experiments for validation of SPECTRA-tube on pooled human sputum. JM designed the *Mtb* qPCR assay. VN conducted experiments for validation of SPECTRA-tube using pooled sputum. AD, SJ, and BJT together wrote the manuscript and prepared figures.

