## Supplementary for "A large-volume sputum dry storage and transportation device for molecular and culture-based diagnosis of tuberculosis"

**Supplementary Figure S1. Melt curve analysis of PCR amplicons for accelerated ageing experiments**

All melt curves show single peaks, confirming that non-specific PCR products were not formed.

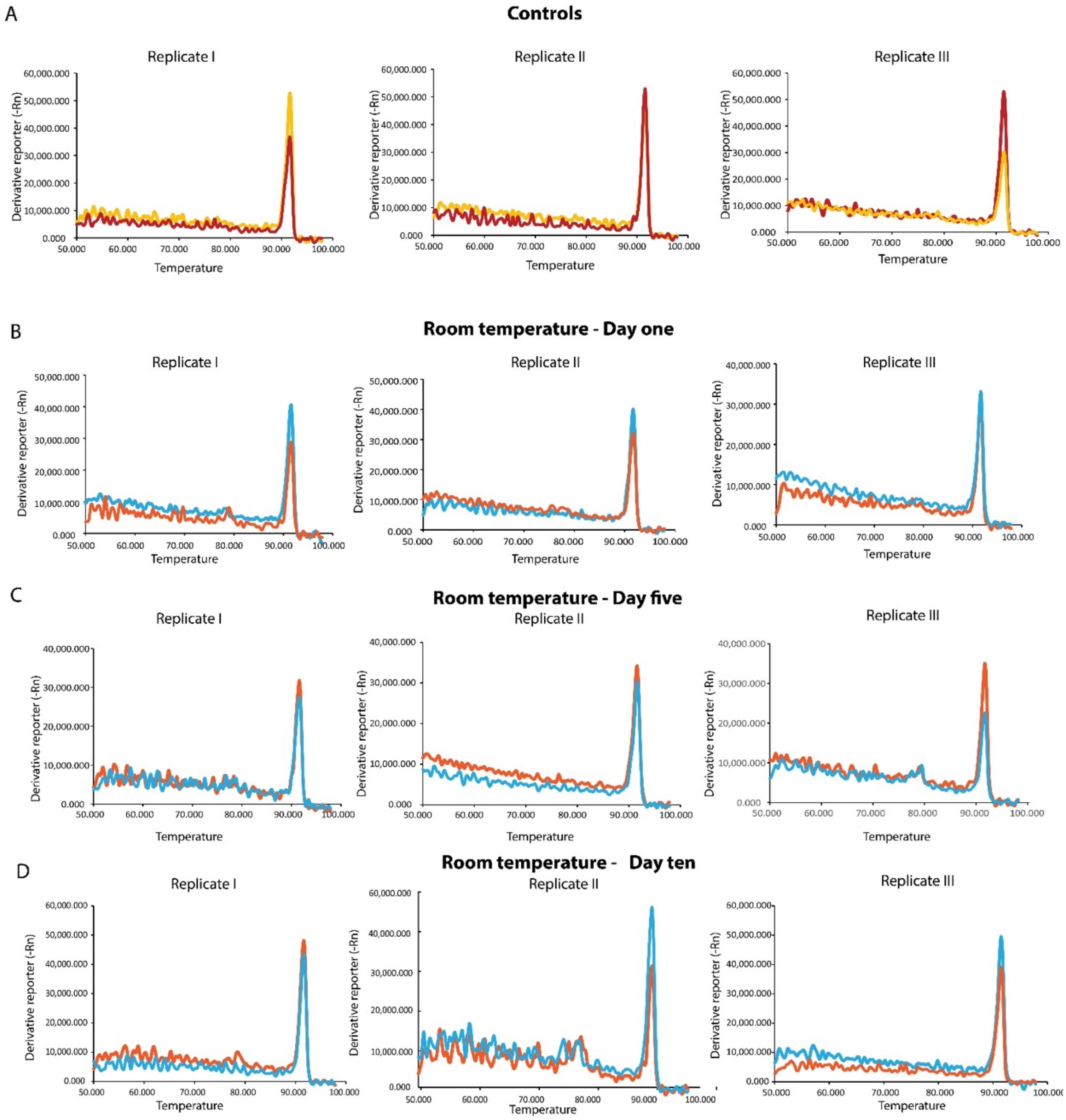

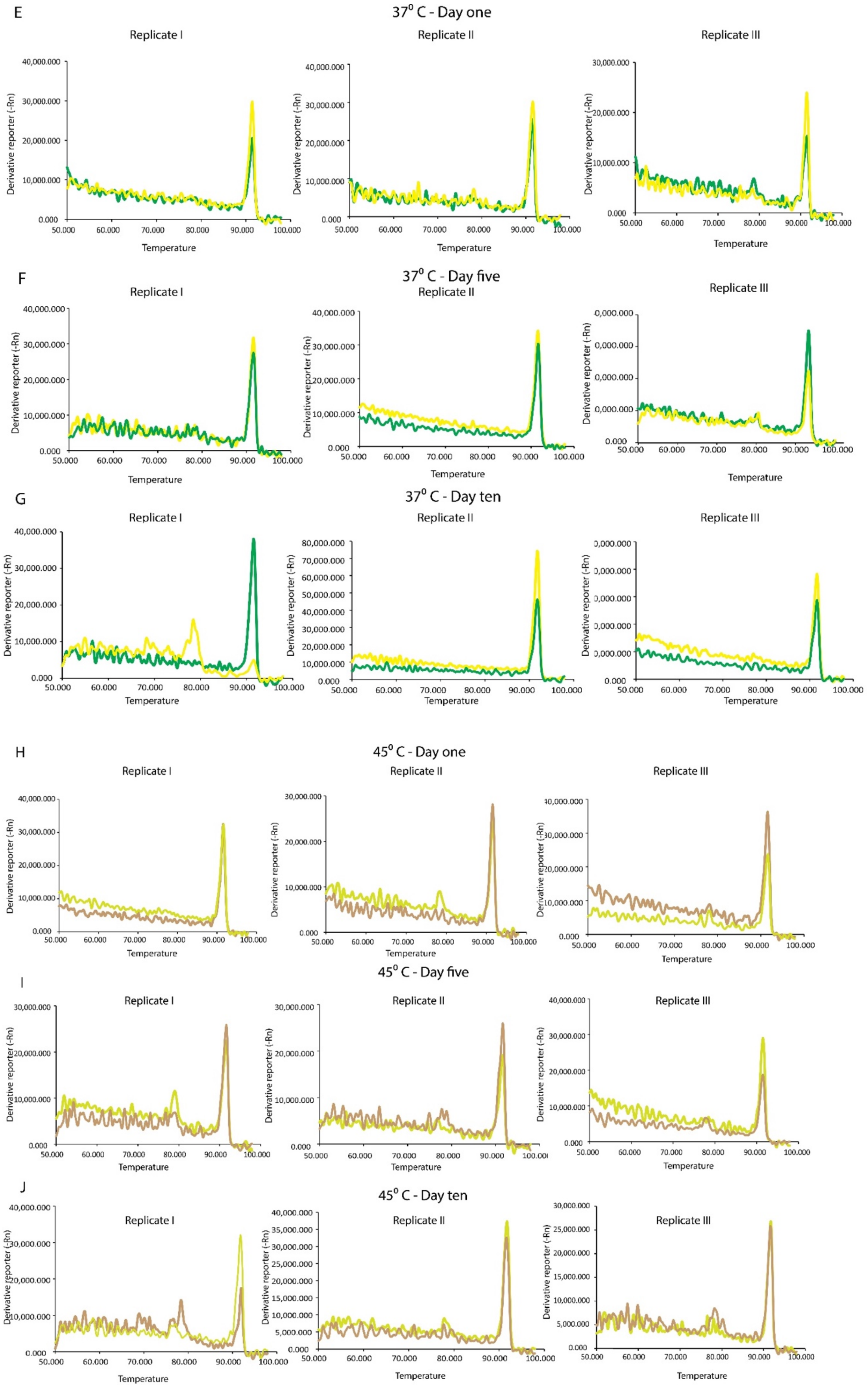
E
37⁰ C - Day one
Replicate I
Replicate II
Replicate III
F
37⁰ C - Day five
Replicate I
Replicate II
Replicate III
G
37⁰ C - Day ten
Replicate I
Replicate II
Replicate III
H
45⁰ C - Day one
Replicate I
Replicate II
Replicate III
I
45⁰ C - Day five
Replicate I
Replicate II
Replicate III
J
45⁰ C - Day ten
Replicate I
Replicate II
Replicate III
Derivative reporter (-Rn)
Temperature

**Figure S1. Melt curve analysis of PCR amplicons for accelerated ageing experiments A. Controls.** Fresh *Msm*-spiked sputum specimens that were not dry-stored. Red, yellow represents reaction duplicates in melt curve analysis. **B,C,D. Room temperature samples.** *Msm*-spiked sputum specimens dry-stored at room temperature and recovered on days 1, 3, and 5. Blue, orange represents reaction duplicates in melt curve analysis. **E,F,G. 37ºC samples.** *Msm*-spiked sputum specimens dry-stored at 37ºC and recovered on days 1, 3, and 5. Yellow, green represents reaction duplicates in melt curve analysis. **H,I,J. 45ºC samples.** *Msm*-spiked sputum specimens dry-stored at 45ºC and recovered on days 1, 3, and 5 respectively. Yellow, brown represents reaction duplicates in melt curve analysis.

**Supplementary Figure S2. Gel electrophoresis analysis of PCR amplicons for accelerated ageing experiments**

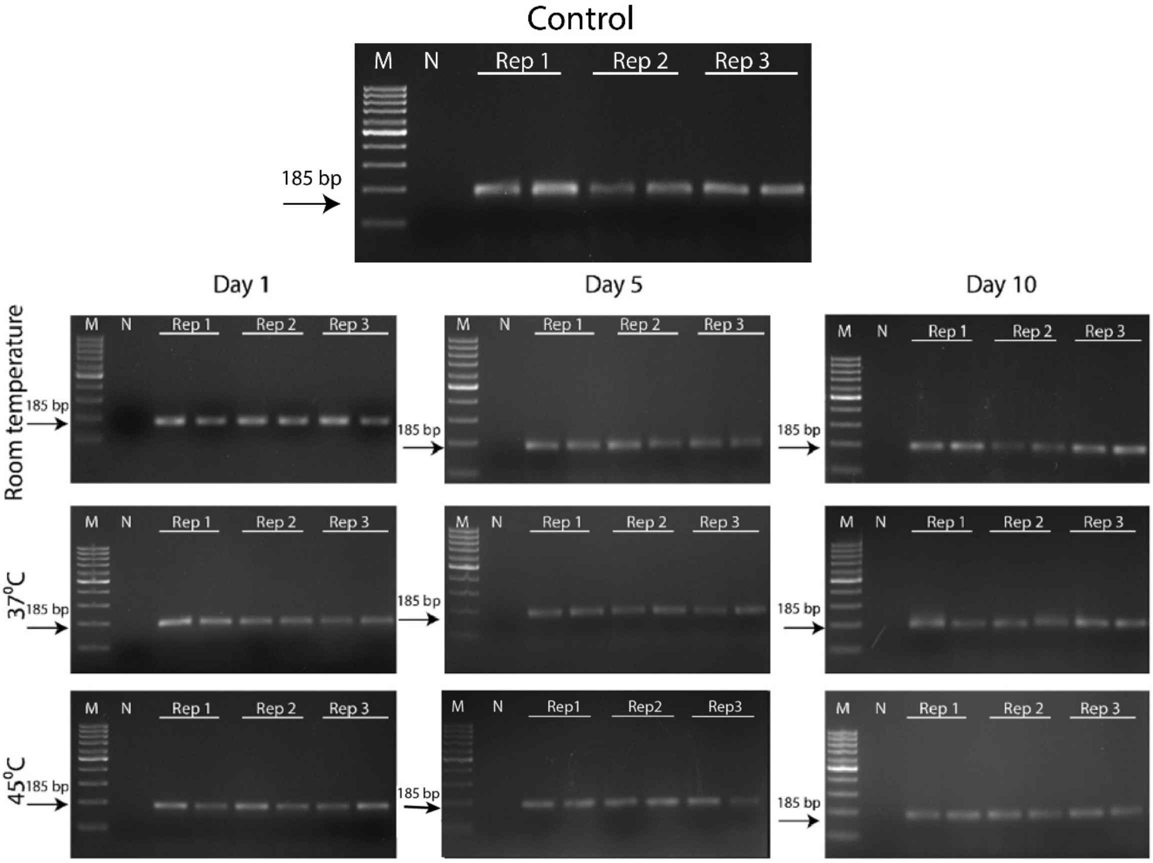


**Figure S2. Gel electrophoresis analysis of PCR amplicons from accelerated ageing studies.** A specific 185bp band is amplified in all the replicates of accelerated aged samples. Control represents a fresh sample that was not dried. M: Marker; N: No template control; Rep1, Rep2, and Rep3: 1cm$^2$ Standard 17 triplicates.

**Supplementary Figure S3. Bacterial culture from *Msm*-spiked mock sputum dried on Standard 17 for 10 days at room temperature, 37ºC, and 45ºC.**

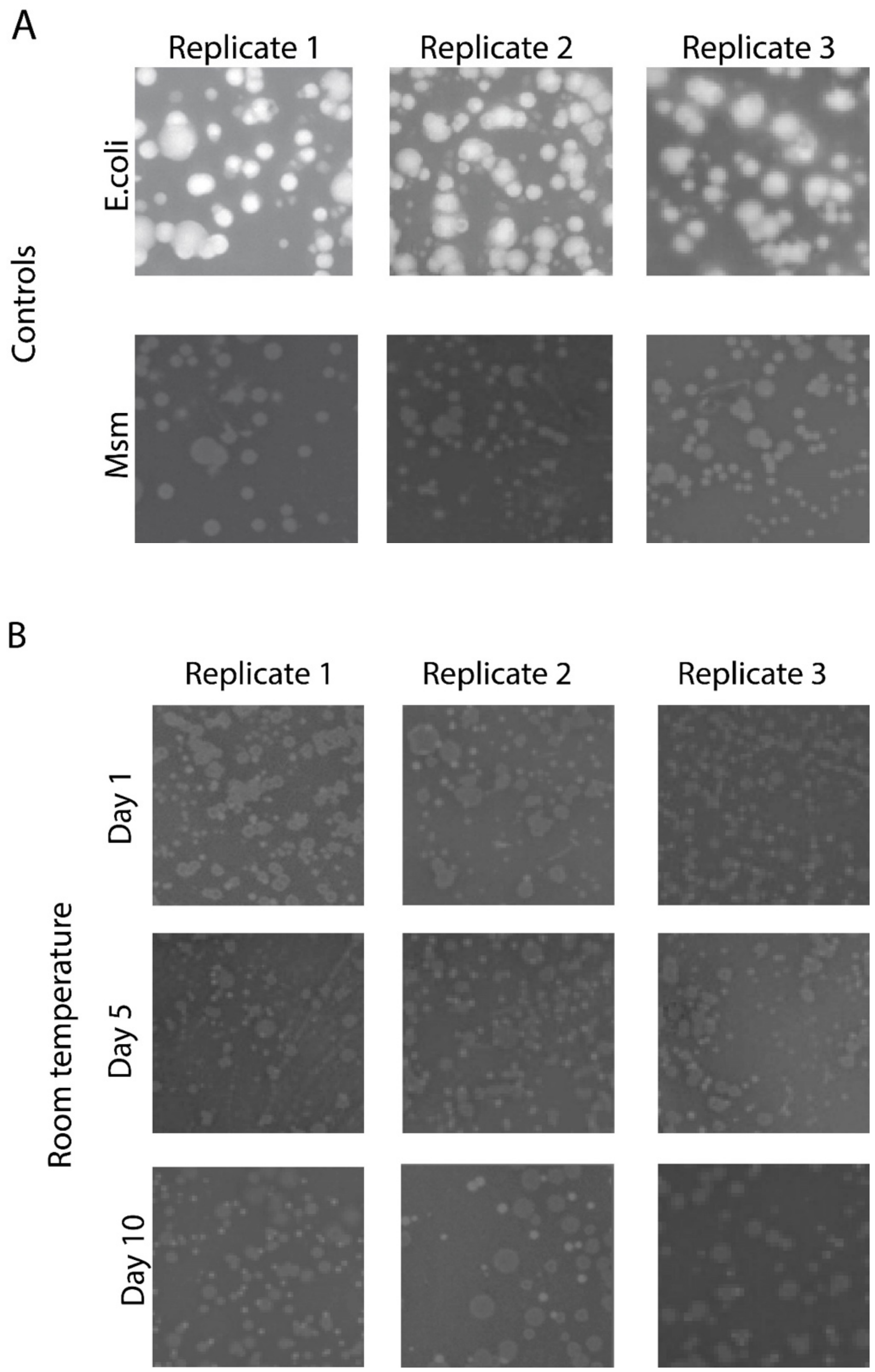

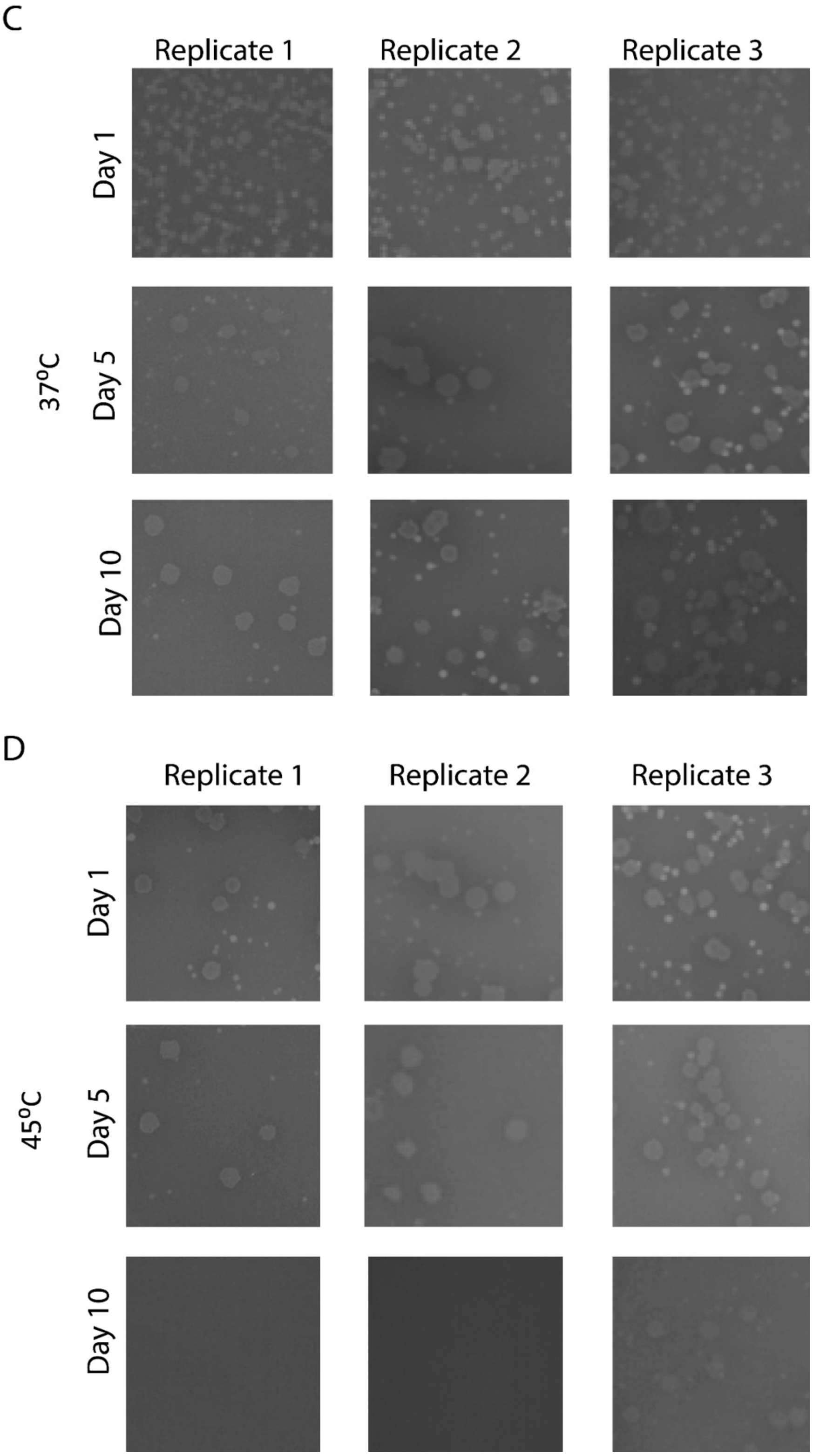


**Figure S3. Bacterial culture from dried mock sputum.** All plates were stained with ethidium bromide (EtBr) and imaged in a UV gel imaging station. **A**: Controls from fresh samples show that *E. coli* bacteria uptake EtBr while *Msm* do not. Lack of EtBr uptake may be used as a test to confirm *Msm* colonies. **B-D**: Bacterial culture from sputums stored at room temperature (B), 37 ºC (C), and 45 ºC (D).

**Supplementary Figure S4. Colony culture PCR from colonies formed from dry stored mock sputum**

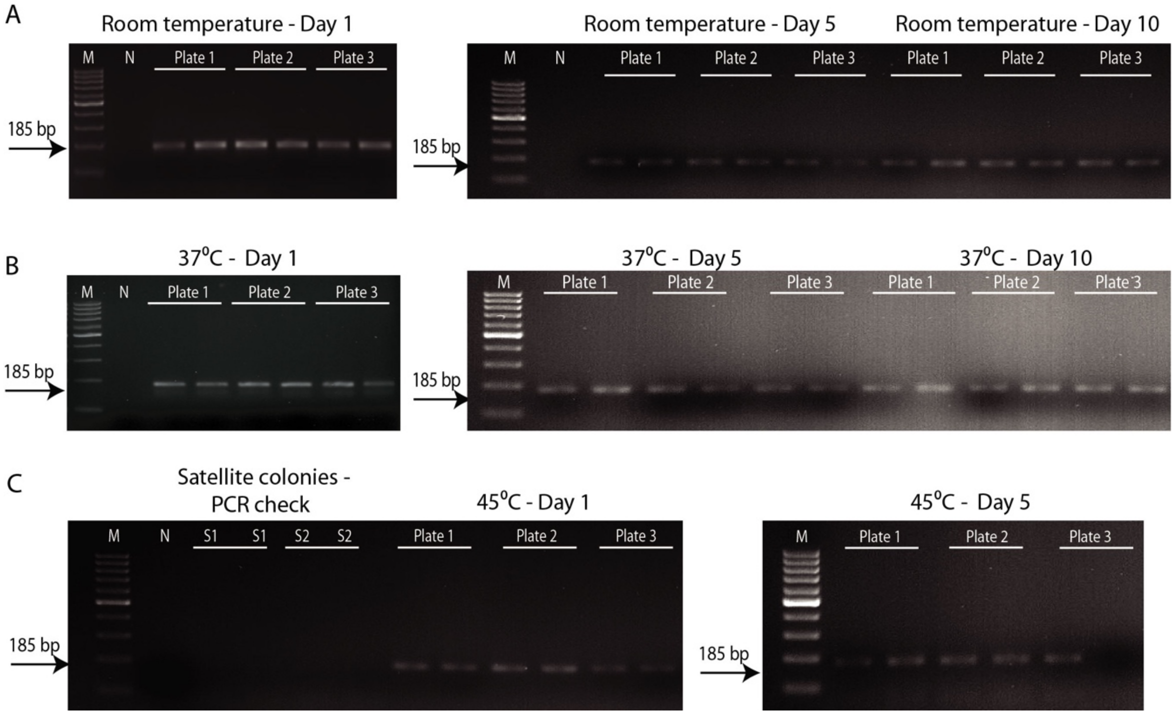


**Figure S4. Colony culture PCR.** Gel electrophoresis analysis of products of PCR from samples stored at room temperature (**A**), 37 ºC (**B**), and 45 ºC (**C**). Presence of the 185 band confirms that the colonies were formed by *Msm* bacteria. A part of panel **C** shows results from colony culture PCR from satellite colonies. Absence of the 185 bp band confirms that the satellite colonies were not formed by *Msm*.

**Supplementary Table S1. Optimization of centrifugation speeds for maximizing sample recovery**

| Sr. No. | Centrifugation Condition | | Efficiency (% weight of added sputum that was recovered) |
|---|---|---|---|
| | RPM | RCF | |
| 1. | 10,000 for 5 min | 16,770 for 5 min | 86.20 |
| 2. | 12,000 for 5 min | 24,149 for 5 min | 90.05 |
| 3. | 6,000 for 3 min then 8,000 for 3 min | 6,037 for 3 min then 10,733 for 3 min | 95.43 |
| 4. | 6,000 for 3 min then 10,000 for 3 min | 6,037 for 3 min then 16,770 for 3 min | 95.71 |

*Assumptions*

- Radius of centrifuge rotor = 15 cm

All experiments were performed using 2-layer SPECTRA-tube devices in which 1.2 mL mock sputum was introduced. Initially only a single RPM for 5 min was used (Sr. No. 1 and 2; Table S1), but efficiency of recovery was only 90.05% at 12,000 RPM. Rather than increasing the RPM further, a 2-stage recovery was tested comprising of a lower RPM for 3 min followed by a higher RPM for 3 min (Sr. No. 3 and 4). This strategy increased recovery to around 95%.

**Supplementary Table S2. Cost of components used in a 2-layer SPECTRA-tube**

| Component | Required quantity | Units | Cost per unit (INR) | Cost in SPECTRA-tube | |
|---|---|---|---|---|---|
| | | | | INR | USD |
| Corning centrifuge tube | 1 | tube | 31 | 31 | 0.42 |
| Standard 17 membrane | 32 | $cm^2$ | 1.09 | 35.12 | 0.24 |
| 2.6 mm thick acrylic | 32 | $cm^2$ | 0.33 | 10.56 | 0.14 |
| Pressure sensitive adhesive | 32 | $cm^2$ | 0.39 | 12.48 | 0.17 |
| PDMS tape | 16 | $cm^2$ | 0.45 | 7.2 | 0.1 |
| 3D printed funnel | 5 | g | 1 | 5 | 0.07 |
| **TOTAL** | | | | **101.36 INR** | **1.38 USD** |

*Assumptions*

- 1 USD (US dollar) = 73.2 INR (Indian Rupee)
